# Metabolic predictors of phenotypic traits can replace and complement measured clinical variables in transcriptome-wide association studies

**DOI:** 10.1101/2022.02.01.478610

**Authors:** Anna Niehues, Daniele Bizzarri, Marcel J.T. Reinders, P. Eline Slagboom, Alain J. van Gool, Erik B. van den Akker, Peter A.C. ’t Hoen, the BBMRI-NL BIOS and Metabolomics Consortia

**Affiliations:** Center for Molecular and Biomolecular Informatics, Radboud Institute for Molecular Life Sciences, Radboud university medical center, Nijmegen, The Netherlands; Translational Metabolic Laboratory, Department Laboratory Medicine, Radboud university medical center, Nijmegen, The Netherlands; Molecular Epidemiology, LUMC, Leiden, The Netherlands; Leiden Computational Biology Center, LUMC, Leiden, The Netherlands; Delft Bioinformatics Lab, TU Delft, Delft, The Netherlands; Max Planck Institute for the Biology of Ageing, Cologne, Germany

## Abstract

Transcriptome-wide association studies (TWAS) can provide valuable insights into biological and disease-underlying mechanisms. For studying clinical effects, availability of (confounding) phenotypic traits is essential. The (re)use of RNA-seq or other omics data can be limited by missing, incomplete, or inaccurate phenotypic information. A possible solution are molecular predictors inferring clinical or behavioral phenotypic traits. Such predictors have been developed based on different omics data types and are being applied in various studies.

In this study, we applied 17 metabolic predictors to infer various traits, including diabetes status or exposure to lipid medication. We evaluated whether these metabolic surrogates can be used as an alternative to reported information for studying the respective phenotypes using TWAS. Our results revealed that in most cases, the use of metabolic surrogates yields similar results compared to using reported information, making them suitable substitutes for such studies.

The application of metabolomics-derived surrogate outcomes opens new possibilities for reuse of multi-omics data sets, especially in situations where availability of clinical metadata is limited. Missing or incomplete information can be complemented by these surrogates, thereby increasing the size of available data sets. Studies would likely also benefit from the use of such surrogates to correct for potential biological confounding. This should be further investigated.

**Author summary:** Transcriptome-wide association studies (TWAS) can be used to associate gene expression levels with phenotypic traits. These associations can provide insights into biological mechanisms including those that underlie diseases. Such studies require molecular profiling data from a large number of individuals as well as information on the phenotypic trait of interest. Biobanks that collect samples and corresponding molecular data, also collect information on phenotypic traits or clinical information. However, this information can be heterogeneous and/or incomplete, or a certain piece of information could be missing entirely. In this study, we apply metabolic predictors to infer various traits, including diabetes status or exposure to lipid medication. We evaluate whether these metabolic surrogates can be used as an alternative to reported information for studying the respective phenotypes using TWAS. Our results reveal that in many cases, the use of metabolic surrogates yields similar results compared to using reported information, making them suitable substitutes for such studies. The possibility of using these surrogate outcomes can thus increase the size of data sets for studies where phenotypic information is incomplete or missing.

## Introduction

Genome-wide association studies (GWAS) have proven to be valuable in uncovering links between genes and a wide range of phenotypic traits. Such findings have lead to the discovery of new disease-related biomarkers and are often the basis for gaining a better understanding of biological processes or disease-mechanisms. Since the introduction of the first GWAS powered by the availability of genome-wide single-nucleotide polymorphism (SNP) profiling, numerous studies have identified thousands of SNP-trait associations [1]. Technological advancements allowing high-throughput profiling of other molecular features, such as transcripts, or DNA methylation sites, also enabled corresponding transcriptome- (TWAS), epigenome- (EWAS), and other ome-wide association studies.

Such studies are susceptible to confounding by biological and technical factors that can influence omics profiles and phenotypic traits of interest. However, measured values to correct for such confounding are often not available. As a solution, differences in cell type composition are commonly accounted for using information contained in the DNA methylation profiles themselves, by either reference-based imputation [2] or reference-free methods such as surrogate variable analysis (methods reviewed in [3] and [4]). Other well-known examples of inferring values for possible confounding factors from DNA methylation profiles include sex [5] and smoking status prediction. Bollepalli et al. [6] trained a smoking status classifier using multinomial LASSO regression. Machine learning approaches have also been applied to other omics data types to predict environmental exposures [7]. Recently, Bizzarri et al. [8] trained predictors on proton NMR-based metabolomics (Nightingale Health) data. The authors applied logistic regression using elastic net regularization to train models for various clinical variables, including physiological measures, environmental exposures, and clinical endpoints. They demonstrated the use of their predicted values as metabolic surrogates to complement missing phenotypic data when performing metabolome-wide association studies (MWAS).

The value of such predictors is not only evident when complementing missing data to account for technical or biological confounding, but also for using them as outcome variables. These molecular surrogates can be used in association studies in order to link molecular features to clinical phenotypes or exposures. Since identified associations in ome-wide association studies often have only moderate effect sizes, a common approach to detect relevant features are cross-cohort meta-analyses [9]. However, the applicability of meta-analyses can be limited by availability of the respective outcomes of interest. Specific clinical, environmental, or phenotypic traits might not be recorded in every cohort, or the data collection might be based on different protocols, making the reported values for these traits not directly comparable.

As more and more multi-omics data sets become available, it becomes possible to make use of molecular predictors to infer phenotypic traits from specific omics layers. We propose their use in subsequent analyses of other omics layers, not only as covariates to account for confounding factors, but also as outcome variables. We here investigated whether values derived from molecular predictors represent a viable alternative to measured or reported clinical or phenotypic traits to serve as outcome variables in TWAS.

To this end, we applied 17 metabolic predictors to metabolomics data from four large population cohorts inferring values for phenotypic, exposure, and clinical traits. We performed TWAS analyses on corresponding RNA-seq data sets employing either reported/measured or inferred values as outcome variables, and systematically compared the respective results of these analyses. For five of the outcomes, where reported values were not available, we evaluated the performance of the metabolic surrogates based on results reported in literature.

## Results

In this study, we performed transcriptome-wide association studies (TWAS) to evaluate the performance of 17 surrogate outcomes that are based on molecular predictors. RNA-seq data used for TWAS and metabolomics data used to infer surrogate outcome values are part of multi-omics data sets of four large Dutch population cohorts: LifeLines (LL) [10], Leiden Longevity Study (LLS) [11], Netherlands Twin Register (NTR) [12], and Rotterdam Study (RS) [13]. An overview of the study workflow is shown in Fig 1. For the different outcomes (Table 1), we compared TWAS results based on metabolic surrogate outcomes to those based on reported outcomes whenever possible. Additionally, results were compared to literature findings.

**Fig 1.**
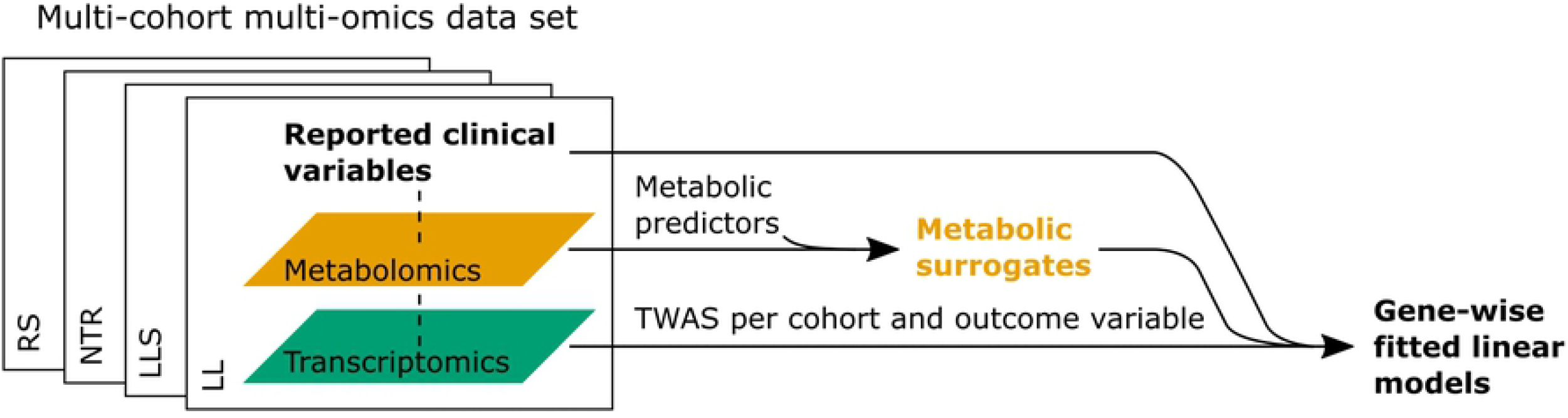
Overview of study workflow. Gene-wise models are fitted for various outcome variables based on reported information or metabolic surrogates, respectively.

**Table 1.** Overview of phenotypic traits.

| Phenotypic trait | Reported outcome | Surrogate outcome |
| --- | --- | --- |
| Low estimated Glomerular Filtration Rate (eGFR) |  | x |
| High triglycerides | x | x |
| High LDL-associated cholesterol | x | x |
| High total cholesterol | x | x |
| Low HDL-associated cholesterol | x | x |
| Diabetes |  | x |
| Metabolic syndrome |  | x |
| Sex | x | x |
| Lipid medication | x | x |
| BMI/obesity status | x | x |
| High high-sensitivity C-reactive protein (hsCRP) | x | x |
| Blood pressure lowering medication |  | x |
| Low hemoglobin | x | x |
| Low white blood cells | x | x |
| Current smoking | x | x |
| Alcohol consumption |  | x |
| High age ($\geq 65$ y.o.) | x | x |

### Metabolic surrogate outcomes

We inferred values for clinical variables by applying molecular predictors to metabolomics data. Recently reported metabolic predictors trained on up to 22 Dutch population cohorts [8] were applied to infer values for outcome variables (Table 1). All 17 metabolic predictors had been shown to perform accurately with mean AUC values > 0.7 in a 5-fold cross-validation approach [8]. In order to avoid emphasis on clinical extremes, the metabolic predictors trained by Bizzarri et al. are based on binary representations using clinical thresholds for continuous variables. The values returned by the predictors are continuous posterior probability scores for belonging to one of the two groups. In most cases, this is the clinical risk group. We here used these predicted values as surrogate outcomes and compared their use in TWAS to reported or measured outcome values.

### Transcriptome-wide association studies

In the next step, the 17 metabolic surrogates were used as outcomes in TWAS. In addition to analyses employing these surrogate outcomes, analyses using measured or reported outcome values were performed. For five outcomes that had limited availability of reported values (eGFR, diabetes, metabolic syndrome, blood pressure lowering medication, and alcohol consumption) only surrogate outcomes were used. In each linear regression model, known biological (age, sex) and technical (flow cell number, white blood cell composition) confounding factors were included (formulas available in S1 Table).

For an initial assessment of the performance of each model, we compared the numbers of significant associations and the effect sizes between outcomes and outcome variable types. Additionally, test statistic (*t*-statistic) bias and inflation were estimated as parameters (mean and standard deviation) of the empirical null distribution using a Bayesian method implemented in the R package `bacon` [26] (Fig 2). Numbers of identified significant associations (Fig 2A) varied strongly across outcomes. Highest numbers were found for the outcomes triglycerides, metabolic syndrome, and white blood cells. For several outcomes, including eGFR, LDL cholesterol, total cholesterol, and alcohol consumption, no or only few significant gene-trait associations were found. For outcomes where TWAS results based on surrogate outcomes could be compared to results for reported variables, the numbers of identified significant associations averaged across all cohorts were higher for the metabolic surrogate outcomes in three cases, and lower in nine cases. However, the variation across the four cohorts was generally higher than the difference between models employing either reported or surrogate outcome variables. Similarly, high variation across cohorts was observed for the other parameters assessed to evaluate the performance of the models. Absolute effect size averaged across all genes (Fig 2B) were generally small, with the highest values observed for the outcomes triglycerides and sex. In 10 cases, the mean absolute effect size averaged across cohorts was lower when using metabolic surrogate outcomes instead of reported variables; in two cases, it was higher. We observed relatively low test statistic bias (Fig 2C) across all outcomes and types of outcome variables. The bias, i.e., the deviation of the empirical null distribution’s mean from zero, averaged across cohorts decreased in four cases, increased in three cases, and remained similar in five cases when employing metabolic surrogate outcomes instead of reported variables. Bias in the RS cohort was often higher than in the other cohorts. This may be explained with the differences in population structure. The RS cohort has a higher average age than the other three cohorts [27], indicating higher bias for the studied clinical variables in older populations. Inflation (deviation of the empirical null distribution’s standard deviation from one, Fig 2D) was highest for the outcome sex. In most cases, inflation averaged across cohorts remained stable when using different outcome variable types. For the outcome total cholesterol, it slightly decreased when using metabolic surrogate outcomes instead of reported variables; for the outcomes lipid medication and hsCRP, it slightly increased.

**Fig 2.**
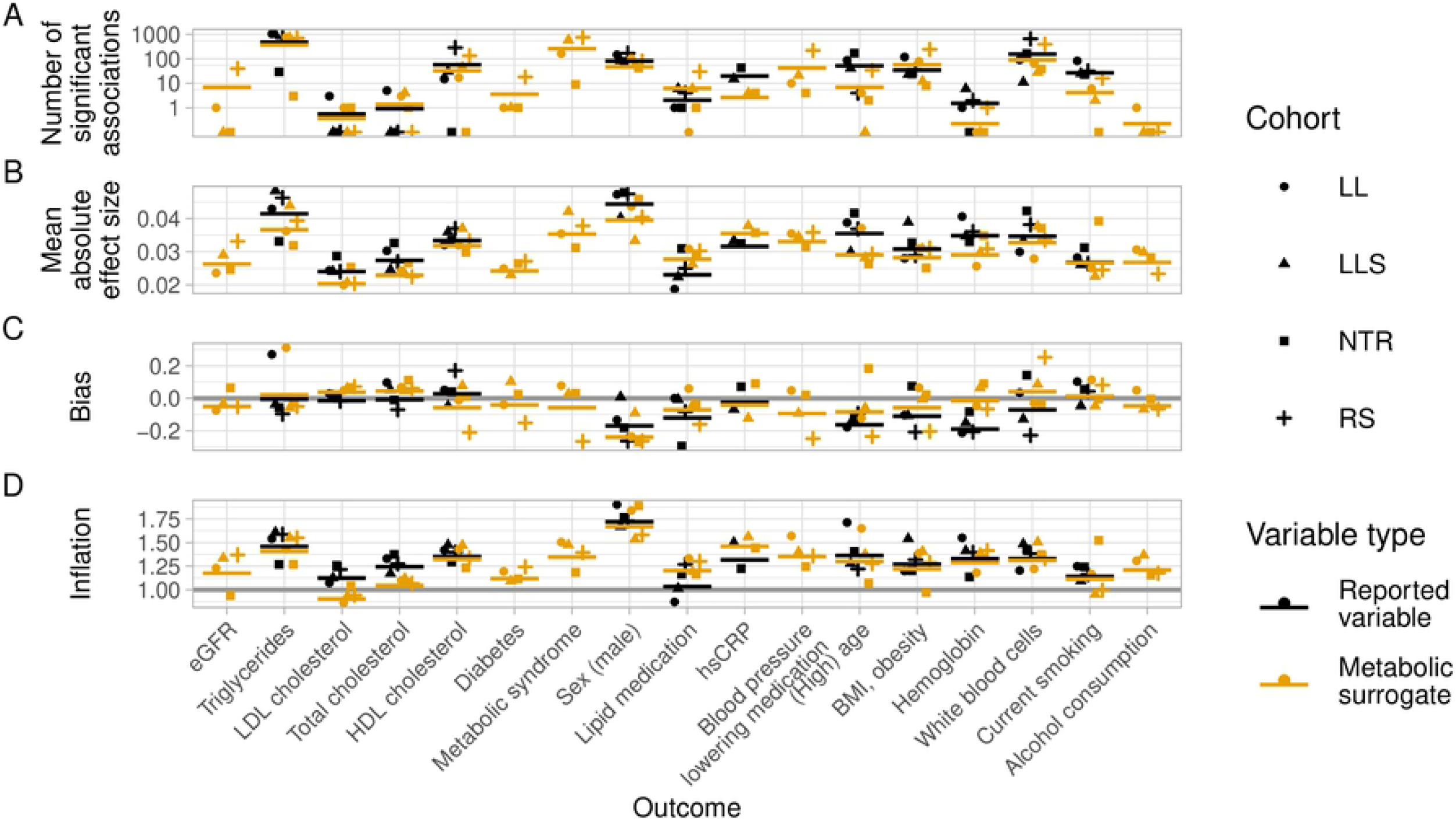
Comparison of TWAS result characteristics. Number of significant associations (based on `bacon`-corrected and Bonferroni-corrected p-values *p_b_adj__* < 0.05) (A), mean absolute effect sizes (based on `bacon`-corrected effect sizes) across all genes (B), and bias (C) and inflation (D) of test statistics (*t*-statistic) for alternative models per comparisons and cohort. Horizontal lines for test-statistic mean = 0 and standard deviation = 1 of theoretical null distribution were added. Comparisons are ordered by performance of metabolic predictors for binary outcome measures. Type of outcome variable is indicated by color: reported or measured variable = black, metabolic surrogate = orange. Mean values across four cohorts (two cohorts for hsCRP) are plotted as horizontal bars. Note the log_10_ scale on the y-axis of the upper plot.

Since the number of significant associations and average effect sizes do not allow to draw conclusions about the similarity of the TWAS results employing different types of variables as outcome, we next performed pairwise comparisons of models with different types of outcome variables, i.e., reported vs. surrogate. Fig 3 shows the correlation of regression coefficients from gene-wise fitted linear models between two different types of outcome variables. Correlation coefficients were generally high for surrogate outcomes based on best-performing metabolic predictors. For outcomes based on predictors with reported AUC > 0.9 (triglycerides, LDL cholesterol, total cholesterol, HDL cholesterol, and sex) [8], the correlation coefficients averaged across cohorts ranged between 0.71 and 0.96. We observed a modest trend for decreasing correlations with decreasing performance of predictors. However, there were exceptions, with sex having highest correlation values, although the predictor’s AUC was reported to be lower than those for triglycerides or cholesterol. Lowest similarity of TWAS results with average absolute Pearson *r* < 0.5 were observed for the outcomes age, white blood cells, smoking, and hemoglobin, the latter having lowest correlation values. We often observed that correlations were lower for the NTR cohort. This could be explained by a technical difference in the metabolic profiles, with NTR missing glutamine [8].

**Fig 3.**
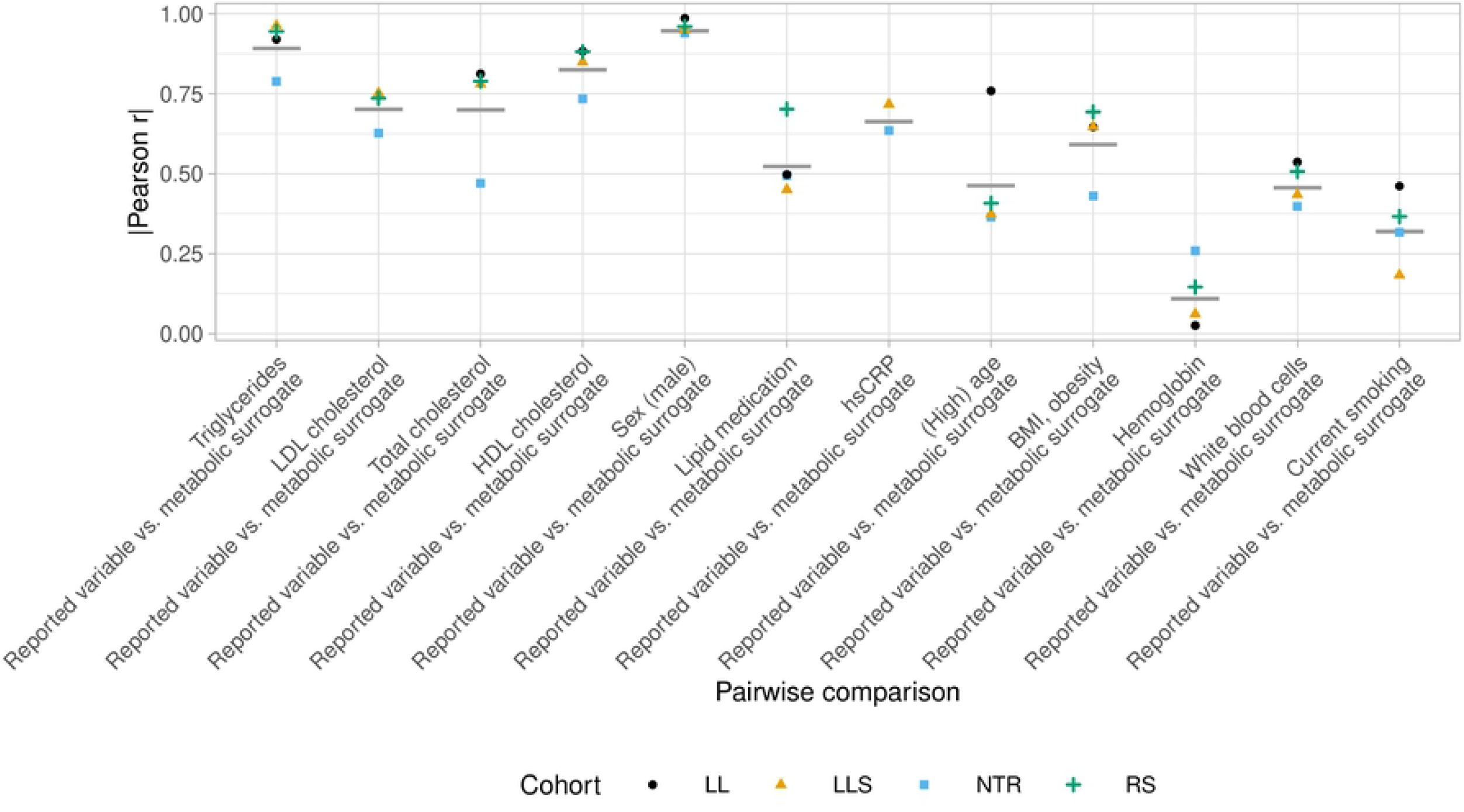
Pairwise comparisons of TWAS results. Absolute Pearson correlation coefficients (Pearson *r*) of `bacon`-adjusted regression coefficients of gene-wise linear models (`limma`/`voom`) for outcome variables in alternative models per comparison and cohort. Comparisons are ordered by performance (AUC) of metabolic predictors for binary outcome measures. Mean values across four cohorts (two cohorts for hsCRP) are plotted as horizontal bars (gray).

### Meta-analyses and replication studies

When comparing two alternative outcome variables, a lower or higher number of found significant associations does not necessarily imply that results are better or worse, since the values alone do not indicate if this is due to a reduction or increase of false positive (noise) or true positive findings, respectively. We observed that the TWAS results differ when surrogate values differ from reported values. However, we do not know which set of results is correct, as reported values could contain inaccuracies. Under the assumption that true positive findings, as opposed to false postive results, can be replicated in different cohorts (validating the results), we performed replication studies to determine which outcome variable type is more consistently reflected in the RNA-seq data. We performed leave-one-cohort-out meta-analyses and replication studies for all comparisons (except for hsCRP where only two cohorts were available) using the approach described by van Rooij et al. [27]. For each comparison, four meta-analyses were performed leaving one cohort out each time, and using the left out cohort for a replication analysis. Fig 4 shows the numbers of significant associations found in each meta-analysis (number of meta-analyzed genes) and the respective percentage of replicated genes. Overall, we did not find substantial differences in the numbers of meta-analyzed genes (Fig 4A) except for the outcomes lipid medication, (high) age, and current smoking. While more genes were meta-analyzed when using the metabolic surrogate for lipid medication, the reported variable yielded more meta-analyzed genes for age and smoking. For the latter two outcomes, results based on metabolic surrogates could not be replicated (Fig 4B) while on average 11-15% of results based on reported outcomes could be replicated. For these two outcomes we had also observed highest differences between number of significations associations (Fig 2A). For other outcome variables, the percentage of replicated genes was quite similar between outcome variable types, but the cohort which was left out for the meta-analysis had a strong impact on the results. Replication rates were generally low with the highest average replaction rate observed for triglycerides with just under 30%. For a number of outcomes, associations could hardly be replicated: LDL-associated cholesterol, hemoglobin, and alcohol consumption. This is in line with the fact that almost no significant associations were found for these outcomes in the individual cohorts (Fig 2).

**Fig 4.**
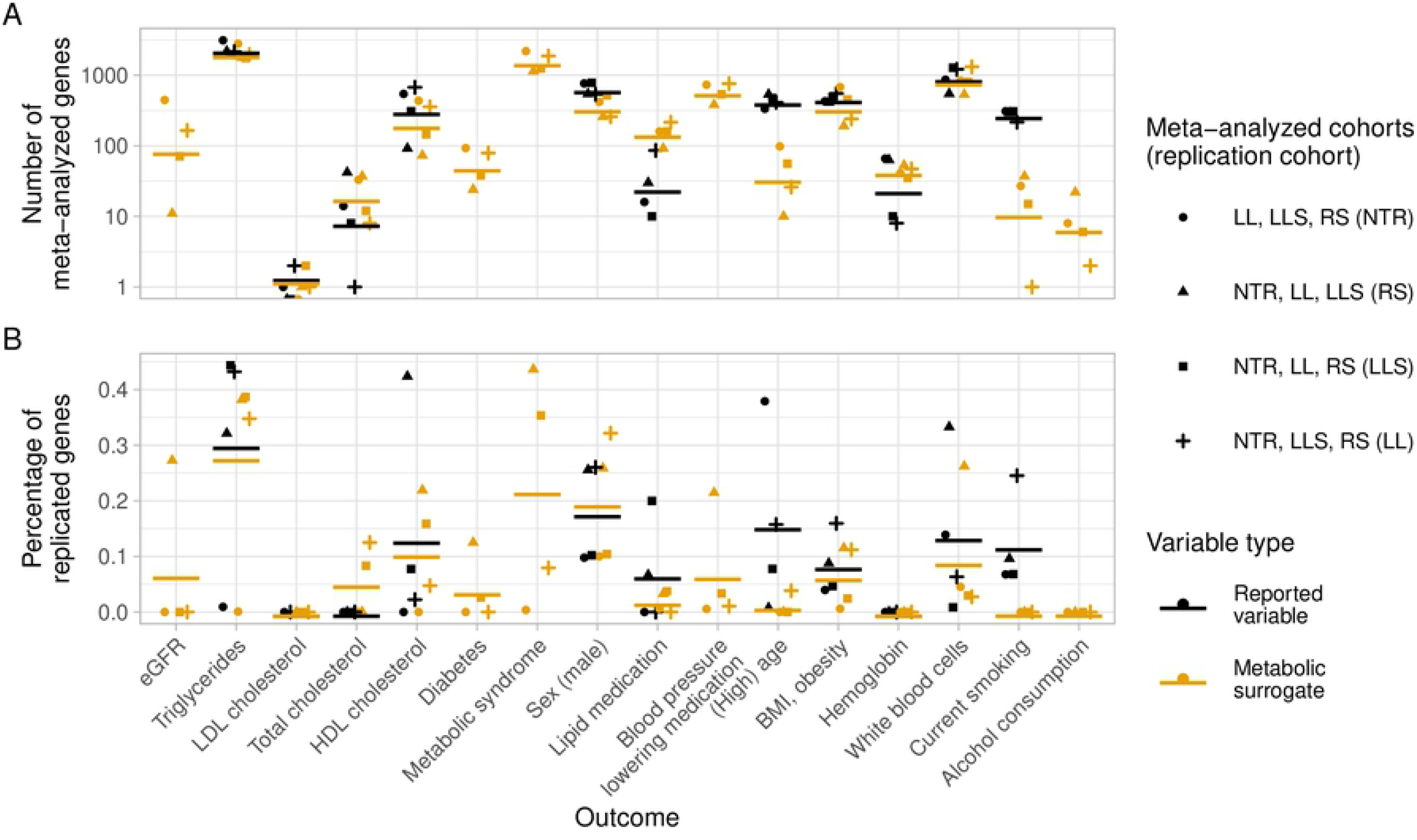
Meta-analyses and replication studies. Number of meta-analyzed genes (significant associations, `bacon`-adjusted *p*-values Bonferroni-adjusted for multiple testing, *p* < 0.05) in leave-one-cohort meta-analyses (A) and percentage of genes replicated (significant associations, Bonferroni-adjusted for multiple testing, *p* < 0.05) in replication cohort (B). Type of outcome variable is indicated by color: reported or measured variable = black, metabolic surrogate = orange. Mean values across four meta-analyses/replication studies are plotted as horizontal bars. Note the log_10_ scale on the y-axis of the upper plot.

### Gene set enrichment analysis

To arrive at a biological interpretation of the TWAS results, we performed gene-set enrichment analyses (GSEA) using pathways from the Reactome database. GSEA was applied to both individual TWAS results and results from a meta-analysis of all four cohorts (Fig 5). For direct comparisons of reported variables and metabolic surrogates, we observed a highest overlap of significantly enriched pathways for HDL cholesterol and triglycerides in all cohorts, with 70-84% (meta-analysis 76%) and 65-80% (meta-analysis 77%) of significantly enriched pathways found by both reported and surrogate outcome, respectively. The overlap for the outcomes total cholesterol, lipid medication, hsCRP, BMI/obesity, and white blood cells was more variable across cohorts. Meta-analyzed results had an overlap of 38-69% and the order of significantly enriched pathways was highly comparable (see S1 Appendix). The results for high age, current smoking, hemoglobin, and LDL cholesterol demonstrated a lower overlap than other outcomes and showed higher variation across the four cohorts compared to other outcomes. This is partially in line with the comparison of gene-wise linear models from TWAS (Fig 3), which showed that results for the outcome hemoglobin based on reported and inferred values were not correlated, and high age and current smoking were only moderately correlated. Since hardly any significant associations were found for hemoglobin (see Fig 2A), the observed signal for this outcome was generally very low in the studied cohorts, independent of the type of outcome variable. It is surprising to observe that almost no significantly enriched pathways were observed for the outcome sex, even though many gene-trait associations were found and could be replicated in the meta-analysis and replication approach.

**Fig 5.**
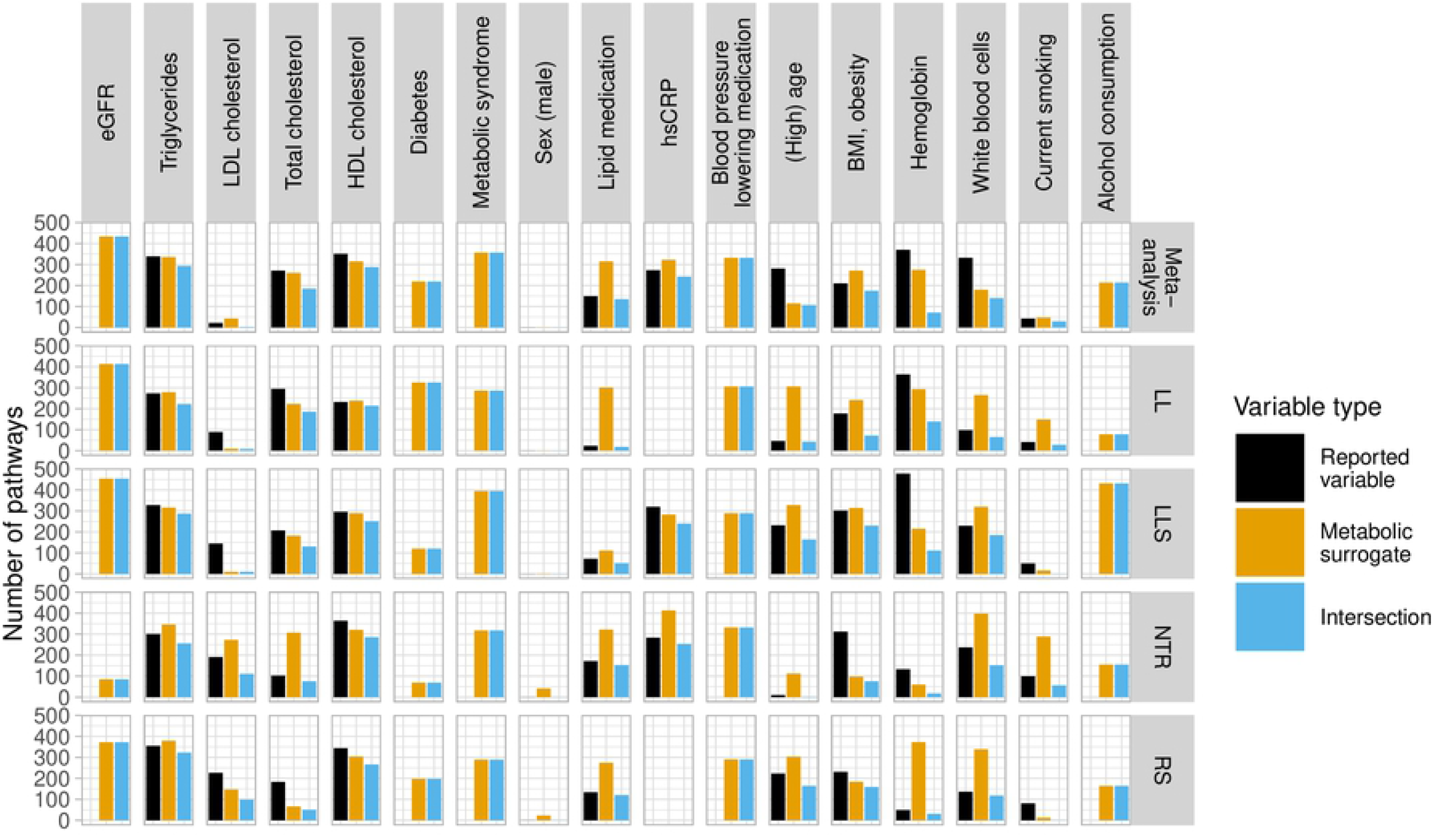
Gene-set enrichment analyses of TWAS results. Numbers of significantly enriched (Bonferroni-adjusted *p* < 0.05) pathways (Reactome) for each outcome found in each cohort and in meta-analysis of all four (two for hsCRP) cohorts (top). Values for each type of outcome variable are represented as colored bars: reported variable = black, metabolic surrogate = orange, intersection, i.e., pathways found by all outcome variables = blue.

In order to evaluate the performance of surrogates for which results could not be compared to results based on reported outcomes, we compared significantly enriched pathways (see S1 Appendix) to the literature. The top-ranked enriched pathways for low eGFR are related to translation. Since low eGFR is an indicator of kidney disease, this is in line with studies reporting increased translational activity to several kidney diseases [30, 31]. For diabetes and metabolic syndrome, which is a risk factor for diabetes, 16 out of the top 20 significantly enriched pathways were found for both outcomes. These include pathways related to translation, signaling, infection, and amino acid deficiency and metabolism. This is in agreement with previously reported results [32, 33]. For alcohol consumption, although almost no significant associations were found in individual TWAS, several significant gene-outcome associations were found when meta-analyzing multiple cohorts (compare Fig 2 and Fig 4). Top-ranked positively enriched Reactome pathways from gene-set enrichment analysis (S1 Appendix), including, e.g., innate immune system, signal transduction, and infectious disease, have been linked previously to chronic alcohol drinking [34].

## Discussion

Our systematic comparison of results of TWAS against outcome variables that were either reported or inferred by molecular predictors generally showed good agreement (Fig 2). Most similar TWAS results across all assessment parameters are those for the outcomes triglycerides and HDL cholesterol. Many significant gene-trait associations were found, of which many could be replicated, and the majority of significantly enriched pathways were found by reported and surrogate outcomes. Regression coefficients of the models including the reported and surrogate outcomes were strongly correlated, and the majority of pathways found by GSEA were obtained by both outcome types. The top-ranked pathways positively enriched for both high triglycerides and low HDL-cholesterol include “GTP hydrolysis and joining of the 60S ribosomal subunit”, “L13a-mediated translational silencing of Ceruloplasmin expression” and “Formation of a pool of free 40S subunits” which participate in the Eukaryotic translation initiation [29] and were previously shown to be enriched in a high-cholesterol and high-fat diet induced hypercholesterolemic rat model [35]. The metabolic predictors for these outcomes are directly related to metabolic markers measured on the Nightingale platform and had shown high performance (AUC ≥ 0.95) [8]. It is expected that results based on predicted outcomes will depend on the accuracy of the prediction. Accordingly, we observed a slightly lower correlation between reported outcomes and metabolic surrogates for molecular predictors that were known to have lower accuracy (Fig 3). In the TWAS for lipid medication and BMI/obesity, the molecular predictors yielded even more significant gene-trait associations than the reported outcomes (Fig 2A) and a higher number of significantly enriched pathways were obtained when using the metabolic surrogates (Fig 5). For lipid medication, this may be related to inaccurate recording of this trait in the questionnaires used. For BMI, this may be explained by the more direct capturing of metabolic processes that are associated with obesity as a combined measure of BMI and waist circumference, because BMI alone is not a perfect indicator of metabolic health [36]. Similarity of results based on surrogate and reported value for sex was high across all assessment parameters, but GSEA did not yield significantly enriched pathways in most cases. It is possible that the genes significantly associated with sex belong to too many different pathways and/or that some genes within a pathway have a positive association while others have a negative association resulting in a failure to identify positively or negatively enriched pathways. For the outcomes total cholesterol and hsCRP, regression coefficients were moderately correlated and GSEA results were very similar. For the outcome white blood cells overlap of significantly enriched pathways in GSEA was smaller. However, the order of top-ranked pathways was similar (see S1 Appendix). The application of metabolic surrogates for outcomes that were not reported in the data and the comparison of TWAS and GSEA results with literature show that these surrogate outcomes allow to transfer information from one data set (the training data) to another, thereby facilitating to study phenotypes or exposures in data sets which would otherwise not be possible.

For the outcomes LDL cholesterol, high age, hemoglobin and current smoking only few gene-trait associations were significantly associated with the outcome, and even fewer in the case of surrogate outcomes. For high age and current smoking the surrogate outcomes also performed worse in the meta-analyses and replication studies compared to reported outcome values. Here, differences between results based on different outcome variable types are reflected in lower correlations of regression coefficients of the gene-wise models and in a smaller overlap in enriched pathways from GSEA. Possible reasons for the differences between surrogates and reported values are lower performance of the metabolic predictors for these outcomes, or a lower biological signal for the respective clinical outcomes and thus increased noise in the studied data. It is known that aging is reflected in transcriptomics data [37], but the metabolic predictor for high age trained on binarized data (≥ 65 y.o.) might not be an ideal surrogate to study this. Alternatively, a metabolomics-based biological age predictor based on continuous data [17] might perform better. Differences between GSEA results of different outcome variable types could also arise from different biological information captured by the metabolic surrogates and by reported or measured values. This phenomenon is known from epigenetic clocks whose age predictions can differ from chronological age, and different clocks can reflect different aspects of biological age [38]). While many of the top-ranked pathways (S1 Appendix) for smoking were found by both outcome variable types, some of the pathways were solely found by using either reported smoking status (“smoking_current”) or metabolic surrogate(“s_current_smoking”). Several pathways related to translation initiation (“Formation of the ternary complex, and subsequently, the 43S complex”, “Translation initiation complex formation”, “Ribosomal scanning and start codon recognition”) were only significantly enriched when using the reported variable as outcome. Translation of mRNA is known to be dysregulated in cancers [39]. Pathways only enriched when using the metabolic surrogate include “Platelet activation, signaling and aggregation” and “Hemostasis”. Increased platelet aggregation has been reported in smokers [40] and platelet-dependent thrombin levels were shown to be increased in smokers and following smoking [41]. This possibly indicates that the reported smoking behavior captures effects of long-term exposure to smoking better, while the metabolic surrogate captures effects of acute smoking. It would be interesting to further investigate which aspects of the clinical phenotypes are captured by the metabolic surrogates. This requires additional phenotypic information. To understand which aspect of smoking behavior is reflected in the omics data current smoking status alone might not be sufficient. More information including pack years and years since smoking cessation could help better understand the information captured by the predictors. It is also possible that different omics types capture different effects better, e.g., short-term and long-term effects. In that case, combining reported outcome variables, and/or molecular surrogates from different omics layers could be very useful, not only to study the effect of a certain exposure, but also to better adjust for confounding factors.

## Conclusion

Based on the systematic comparison of TWAS results using either reported variables as outcome or surrogate outcomes inferred from another omics profile, we conclude that metabolic surrogates yield similar results as reported variables. The availability of these surrogate outcomes extends the possibilities of studying various clinical outcomes in population cohorts. It can enable the reuse of existing multi-omics data with limited reported clinical (meta)data. This allows for inclusion of more cohorts in meta-analyses, even when outcomes of interest were not reported for all cohorts. This approach also increases possibilities to study clinical outcomes by allowing to infer important confounding factors. Especially investigations that rely on reuse of existing data, .e.g. in the case of rare disease studies which often also suffer from low sample sizes, will benefit from this approach.

## Methods

### Data

In this study, we analyzed RNA-seq and metabolomics data from four large Dutch population cohorts: LifeLines (LL) [10], Leiden Longevity Study (LLS) [11], Netherlands Twin Register (NTR) [12], and Rotterdam Study (RS) [13, 14]. The data is provided by the Dutch node of the European Biobanking and BioMolecular Resources and Research Infrastructure (BBMRI-NL).

RNA-seq data was generated by the BBMRI-NL Biobank-based Integrative Omics Study (BIOS) Consortium at the Human Genotyping facility (HugeF) of ErasmusMC, the Netherlands. RNA sample processing and sequencing is described in detail by Zhernakova et al. [15]. Briefly, total RNA was extracted from whole blood, depleted of globin transcripts, and paired-end sequencing of 2×50-bp reads was conducted using the Illumina HiSeq 2000 platform. Read alignment to reference genome hg19 was performed using STAR (v2.3.0). We used the “Freeze2 unrelated data sets”, which contain maximum sets of unrelated individuals and are available within the BIOS workspace at the SURF Research Cloud via the R package `BBMRIomics` v3.4.2 [16].

Metabolomics data was generated by the BBMRI-NL Metabolomics Consortium in 2014 as described by van den Akker et al. [17]. Briefly, metabolite concentrations were measured in EDTA plasma by proton nuclear magnetic resonance (^1^H-NMR) spectroscopy on the platform of the Nightingale Health Group (Helsinki, Finland) [18].

### Ethics statement

Written informed consent was obtained previously from all participants of the LL, LLS, NTR and RS biobanks in accordance with the ethical and institutional regulations. The LL study was approved by the ethics committee of the University Medical Centre Groningen, document no. METC UMCG LLDEEP: M12.113965. All participants signed an informed consent form prior to study enrolment [10]. The Leiden Longevity Study was approved by the Medical Ethical Committee of the Leiden University Medical Center (METC LDD) and informed consent was obtained from all subjects. All procedures performed in studies involving human participants were in accordance with the ethical standards of the institutional and/or national research committee and with the 1964 Helsinki declaration and its later amendments or comparable ethical standards [11]. The Netherlands Twin Register [12] study protocol was approved by the Central Ethics Committee on Research Involving Human Subjects of the VU University Medical Centre, Amsterdam, an Institutional Review Board registered with the U.S. Office of Human Research Protections (Institutional Review Board no. IRB00002991, Federal-wide Assurance no. FWA00017598). The study is registered with the Dutch Data Protection Authority (no. m1412317). In accordance with the Declaration of Helsinki, the Netherlands Twin Register obtained informed consent from all participants prior to their entering the study. The Rotterdam Study has been approved by the institutional review board (Medical Ethics Committee) of the Erasmus Medical Center (registration number MEC 02.1015) and by the Dutch Ministry of Health, Welfare and Sport (Population Screening Act WBO, license number 1071272-159521-PG). The Rotterdam Study Personal Registration Data collection is filed with the Erasmus MC Data Protection Officer under registration number EMC1712001. The Rotterdam Study has been entered into the Netherlands National Trial Register (NTR; https://www.trialregister.nl/) under catalogue number Trial NL6645 (NTR6831). All participants provided written informed consent to participate in the study and to have their information obtained from treating physicians [14, 21].

### Data analysis

All analyses were implemented in an R v4.0.3 [19] workflow employing R packages `renv` v0.14.0 for package management and `drake` v7.13.2 for workflow management. The analyses were run in the BIOS workspace of the SURF Research Cloud which is part of the multi-omics analysis platform of BBMRI-NL. The code to run the analyses in available in GitHub (https://github.com/niehues/bbmri_surrogates_twas) and archived in Zenodo [20].

### Data preprocessing

#### Normalization of values for clinical traits

Numeric values were used for all reported clinical variables. In case of categorical variables, they were binarized as follows. For smoking status, “current smoker” was coded as 1, and “former-smoker” and “non-smoker” were coded as 0; for sex, “male” and “female” were coded as 1 and 0, respectively; for lipid medication, “statins” were coded as 1, and “no” and “yes, but no statins” were coded as 0. In order to be able to compare effect sizes in TWAS, all clinical variables were standardized to zero-mean and unit-variance (z-score normalization).

#### RNA-seq data preprocessing

Samples with more than 10% missing values in the RNA-seq data were excluded from the analysis. Additionally, for comparisons of models employing either reported or inferred values as outcome, samples missing reported values were excluded. Subsequently, features (transcripts) missing in more than 10% of the samples were removed from the data set. Number of retained samples and features are given in S1 Table.

The RNA-seq read counts as provided by the `BBMRIomics` R package were then normalized and transformed as follows. Scaling factors for library sizes were calculated using the trimmed mean of M-values (TMM) methods [22] implemented in the R/Bioconductor package `edgeR` v3.32.1 [23]. Using these scaling factors to adjust for sequencing depth, counts were transformed to log_2_ counts-per-million (CPM) reads, their mean-variance relationships were estimated using `voom` [24] implemented in the R/Bioconductor package `limma` v3.46.0, and the associated observation-level weights were used in the subsequent linear modeling to adjust for heteroscedasticity.

#### Metabolomics data preprocessing

The Nightingale Health metabolomics features inquired are the 56 variables selected by van den Akker et al. [17]. Outliers were identified as the samples having more than 1 missing observation (301 removed), more than 1 datapoint under the detection limit (49 removed) and having a value more than 5 standard deviations away from the overall mean observed within BBMRI-NL (0 removed). Remaining with a total of 12926 samples (LLS = 2343, LL = 1475, RS = 5136, NTR = 3972). The remaining 4210 missing values (0.58% of the entire dataset) were imputed as zero and then we z-scaled the metabolic features using the mean and standard deviations observed in BBMRI-NL. The samples were further filtered based on availability of corresponding RNA-seq data. Sample sizes per cohort and model are listed in S1 Table.

### Metabolic surrogates

Seventeen logistic regression models trained on up to 22 cohorts, including LL and RS, were applied to the dataset as described in Bizzarri et al. [8]. The surrogates used in this study are the posterior probabilities which represent how likely an individual is at risk for each of the inquired common clinical variables.

### Transcriptome-wide association studies (TWAS)

Gene-wise linear regression models were fitted using `limma` [25]. Known potential biological (sex, age) and technical confounders (flow cell number, white blood cell composition) were included in the models. TWAS was performed separately for the respective type of outcome variable, i.e., reported variable or metabolic surrogate. Parameters for each TWAS linear model are summarized in S1 Table. We adjusted *p*-values and effect sizes for statistical bias and inflation using the Bayesian method bacon [26], which estimates bias and inflation as parameters from the empirical null distribution of test statistics (*t*-statistic). Additionally, *p*-values were adjusted for multiple testing using the Bonferroni method. TWAS results for two different variable types for the same outcome, were compared by calculating Pearson correlation coefficients of regression coefficients from gene-wise fitted models.

### Meta-analysis

Leave-one-cohort-out meta-analyses and replication studies in left out cohort were performed as described in [27].

### GSEA

Gene-set enrichment analyses (GSEA) were performed using the R/Bioconductor package `fgsea` v1.16.0 [28] and gene sets retrieved from the Reactome Pathway Database [29]. Genes were ranked by – log_10_(*p_b_*) * |*β_b_*| with *p_b_* = `bacon`-corrected p-value and *beta_b_* = `bacon`-corrected effect size. The number of permutations for initial estimation of p-values was set to 1 × 10^4^; the boundary for calculating p-values was set to 1 × 10^-50^.

## Supporting information

**S1 Table. TWAS model parameters.** Variable names, sample and feature numbers, and covariates per TWAS model and cohort.

**S1 Appendix. Significantly enriched pathways from GSEA.** S1 Appendix contains plots showing significantly enriched pathways for each outcome variable. GSEAs are based on meta-analyzed TWAS results.

## Acknowledgments

This research was financially supported by EATRIS-Plus (https://eatris.eu/projects/eatris-plus/), which has received funding from the European Union’s Horizon 2020 research and innovation programme under grant agreement no. 871096, The Netherlands X-omics Initiative (https://x-omics.nl/), which is (partly) funded by the Dutch Research Council (NWO), project no. 184.034.019, and by the Biobank-based Integrative Omics Study (BIOS) Consortium (https://www.bbmri.nl/acquisition-use-analyze/bios) and the BBMRI Metabolomics Consortium (https://www.bbmri.nl/Omics-metabolomics) which are funded by BBMRI-NL, a Research Infrastructure financed by NWO, project nos. 184.021.007 and 184033111. We thank the biobanks and participants of LifeLines (https://www.lifelines.nl/), the Leiden Longevity Study (https://leidenlangleven.nl/), the Netherlands Twin Register (https://tweelingenregister.vu.nl/), and the Rotterdam Study (http://www.epib.nl/research/ergo.htm). Special thanks also to Leon Mei, Davy Cats, Martin Brandt and the SURF RSC Team for their support and management of data and computational infrastructure.

## Members of the BBMRI-NL BIOS Consortium

### Management Team

Bastiaan T. Heijmans (chair)^1^, Peter A.C. ‘t Hoen^2^, Joyce van Meurs^3^, Rick Jansen^5^, Lude Franke^6^.

### Cohort collection

Dorret I. Boomsma^7^, René Pool^7^, Jenny van Dongen^7^, Jouke J. Hottenga^7^ (Netherlands Twin Register); Marleen M.J. van Greevenbroek^8^, Coen D.A. Stehouwer^8^, Carla J.H. van der Kallen^8^, Casper G. Schalkwijk^8^ (Cohort study on Diabetes and Atherosclerosis Maastricht); Cisca Wijmenga^6^, Lude Franke^6^, Sasha Zhernakova^6^, Ettje F. Tigchelaar^6^ (LifeLines Deep); P. Eline Slagboom^1^, Marian Beekman^1^, Joris Deelen^1^, Diana van Heemst^9^ (Leiden Longevity Study); Jan H. Veldink^10^, Leonard H. van den Berg^10^ (Prospective ALS Study Netherlands); Cornelia M. van Duijn^4^, Bert A. Hofman^11^, Aaron Isaacs^4^, André G. Uitterlinden^3^ (Rotterdam Study).

### Data generation

Joyce van Meurs (Chair)^3^, P. Mila Jhamai^3^, Michael Verbiest^3^, H. Eka D. Suchiman^1^, Marijn Verkerk^3^, Ruud van der Breggen^1^, Jeroen van Rooij^3^, Nico Lakenberg^1^.

### Data management and computational infrastructure

Hailiang Mei (Chair)^12^, Maarten van Iterson^1^, Michiel van Galen^2^, Jan Bot^13^, Dasha V. Zhernakova^6^, Rick Jansen^5^, Peter van’t Hof^12^, Patrick Deelen^6^, Irene Nooren^13^, Peter A.C. ‘t Hoen^2^, Bastiaan T. Heijmans^1^, Matthijs Moed^1^.

### Data analysis group

Lude Franke (Co-Chair)^6^, Martijn Vermaat^2^, Dasha V. Zhernakova^6^, René Luijk^1^, Marc Jan Bonder^6^, Maarten van Iterson^1^, Patrick Deelen^6^, Freerk van Dijk^14^, Michiel van Galen^2^, Wibowo Arindrarto^12^, Szymon M. Kielbasa^15^, Morris A. Swertz^14^, Erik. W van Zwet^15^, Rick Jansen^5^, Peter-Bram ‘t Hoen (Co-Chair)^2^, Bastiaan T. Heijmans (Co-Chair)^1^.

### Affiliations of BIOS members

**1** Molecular Epidemiology, Department of Biomedical Data Sciences, Leiden University Medical Center, Leiden, The Netherlands

**2** Department of Human Genetics, Leiden University Medical Center, Leiden, The Netherlands

**3** Department of Internal Medicine, ErasmusMC, Rotterdam, The Netherlands

**4** Department of Genetic Epidemiology, ErasmusMC, Rotterdam, The Netherlands

**5** Department of Psychiatry, VU University Medical Center, Neuroscience Campus Amsterdam, Amsterdam, The Netherlands

**6** Department of Genetics, University of Groningen, University Medical Centre Groningen, Groningen, The Netherlands

**7** Department of Biological Psychology, VU University Amsterdam, Neuroscience Campus Amsterdam, Amsterdam, The Netherlands

**8** Department of Internal Medicine and School for Cardiovascular Diseases (CARIM), Maastricht University Medical Center, Maastricht, The Netherlands

**9** Department of Gerontology and Geriatrics, Leiden University Medical Center, Leiden, The Netherlands

**10** Department of Neurology, Brain Center Rudolf Magnus, University Medical Center Utrecht, Utrecht, The Netherlands

**11** Department of Epidemiology, ErasmusMC, Rotterdam, The Netherlands

**12** Sequence Analysis Support Core, Department of Biomedical Data Sciences, Leiden University Medical Center, Leiden, The Netherlands

**13** SURFsara, Amsterdam, the Netherlands

**14** Genomics Coordination Center, University Medical Center Groningen, University of Groningen, Groningen, the Netherlands

**15** Medical Statistics, Department of Biomedical Data Sciences, Leiden University Medical Center, Leiden, The Netherlands

## Members of the BBMRI-NL Metabolomics Consortium

### Cohort collection

J.M. Geleijnse^1^, E. Boersma^2^, W.E. van Spil^3^, M.M.J. van Greevenbroek^4,5^, C.D.A. Stehouwer^4,5^, C.J.H. van der Kallen^4,5^, I.C.W. Arts^5,6,7^, F. Rutters^8,9^, J.W.J. Beulens^8,9^, M. Muilwijk^8,10^, P.J.M. Elders^8,10^, L.M. ‘t Hart^8,9,11,12^, M. Ghanbari^13,14^, M.A. Ikram^13^, M.G. Netea^15^, M. Kloppenburg^16,17^, Y.F.M. Ramos^18^, N. Bomer^19^, I. Meulenbelt^18^, K. Stronks^20^, M.B. Snijder^20^, A.H. Zwinderman^21^, B.T. Heijmans^18^, L.H. Lumey^22^, C. Wijmenga^23^, J. Fu^23,24^, A. Zhernakova^23^, J. Deelen^25,18^, S.P. Mooijaart^26^, M. Beekman^18^, P.E. Slagboom^18,25^, G.L.J. Onderwater^27^, A.M.J.M. van den Maagdenberg^28,27^, G.M. Terwindt^27^, C.Thesing^29,8^, M. Bot^29,8^, B.W.J.H. Penninx^29,8^, S. Trompet^30,26^, J.W. Jukema^30^, N. Sattar^31^, I.C.C. van der Horst^32^, P. van der Harst^33^, C. So-Osman^34,35^, J.A. van Hilten^36^, R.G.H.H. Nelissen^37^, I.E. Höfer^38^, F.W. Asselbergs^39,40^, P. Scheltens^41^, C.E. Teunissen^42^, W.M. van der Flier^43,41^, J. van Dongen^29,8^, R. Pool^29^, A.H.M. Willemsen^29,8^, D.I. Boomsma^29,8^.

### Sample logistics, database and catalogue

H.E.D. Suchiman^18^, J.J.H. Barkey Wolf^18^, M. Beekman^18^, D. Cats^45^, H. Mei^45^, M. Slofstra^23^, M. Swertz^46,23^, M.J.T. Reinders^47,48^, E.B. van den Akker^47,18^.

### Steering committee

D.I. Boomsma^29,8^, M.A. Ikram^13^, P.E. Slagboom^18,25^.

### Affiliations of BBMRI-NL Metabolomics members

**1** Division of Human Nutrition and Health, Wageningen University, Wageningen, The Netherlands

**2** Thorax centre, Erasmus Medical Centre, Rotterdam, the Netherlands

**3** Department of Rheumatology & Clinical Immunology, University Medical Center Utrecht, Utrecht, The Netherlands

**4** Department of Internal Medicine, Maastricht University Medical Center (MUMC+), Maastricht, The Netherlands

**5** School for Cardiovascular Diseases (CARIM), Maastricht University, Maastricht, the Netherlands

**6** Department of Epidemiology, Maastricht University, Maastricht, the Netherlands

**7** Maastricht Center for Systems Biology, Maastricht University, Maastricht, the Netherlands

**8** Amsterdam Public Health Research Institute, Amsterdam, The Netherlands

**9** Department of Epidemiology and Biostatistics, Amsterdam University Medical Center, Vrije Universiteit, Amsterdam, the Netherlands

**10** Department of General Practice and Elderly Care Medicine, Amsterdam University Medical Center, Vrije Universiteit, Amsterdam, the Netherlands

**11** Department of Epidemiology and Biostatistics, Amsterdam University Medical Center, Vrije Universiteit, Amsterdam, the Netherlands

**12** Department of Cell and Chemical Biology, Leiden University Medical Center, Leiden, the Netherlands

**13** Department of Epidemiology, Erasmus MC, University Medical Center, Rotterdam, The Netherlands

**14** Department of Genetics, School of Medicine,, Mashhad University of Medical Sciences, Mashhad, Iran

**15** Department of Internal Medicine and Radboud Center for Infectious Diseases, Radboud University Medical Center, Nijmegen, The Netherlands

**16** Department of Clinical Epidemiology, Leiden University Medical Centre, Leiden, The Netherlands

**17** Department of Rheumatology, Leiden University Medical Center, The Netherlands

**18** Department of Biomedical Data Sciences, Section of Molecular Epidemiology, Leiden University Medical Center, Leiden, The Netherlands

**19** Department of Experimental Cardiology, University of Groningen, University Medical Center Groningen, Groningen, The Netherlands

**20** Department of Public Health, Academic Medical Center, University of Amsterdam, Amsterdam, The Netherlands

**21** Department of Clinical Epidemiology, Biostatistics, and Bioinformatics, Academic Medical Centre, University of Amsterdam, Amsterdam, The Netherlands

**22** Department of Epidemiology, Mailman School of Public Health, Columbia University, New York, NY 10032

**23** Department of Genetics, University Medical Center Groningen, Groningen, The Netherlands

**24** Department of Pediatrics, University Medical Center Groningen, Groningen, The Netherlands

**25** Max Planck Institute for Biology of Ageing, Cologne, Germany

**26** Department of Internal Medicine, Division of Gerontology and Geriatrics, Leiden University Medical Centre, Leiden, The Netherlands

**27** Department of Neurology, Leiden University Medical Center, Leiden, The Netherlands

**28** Department of Human Genetics, Leiden University Medical Center, Leiden, The Netherlands

**29** Department of Biological Psychology, Amsterdam University Medical Center, Vrije Universiteit, Amsterdam, The Netherlands

**30** Department of Cardiology, Leiden University Medical Center, Leiden, The Netherlands

**31** Institute of Cardiovascular and Medical Sciences, Cardiovascular Research Centre, University of Glasgow, Glasgow, UK

**32** Department of Critical Care, University Medical Center Groningen, Groningen, The Netherlands

**33** Department of Cardiology, University Medical Center Utrecht, Utrecht, The Netherlands

**34** Sanquin Blood Bank, Leiden and Department of Haematology, Groene Hart Hospital, Gouda, the Netherlands

**35** International Society of Blood Transfusion (ISBT), Amsterdam, The Netherlands

**36** Unit of Transfusion Medicine, Sanquin Blood Bank, Leiden, The Netherlands

**37** Department of Orthopaedics, Leiden University Medical Center, Leiden, The Netherlands

**38** Department of Clinical Chemistry and Hematology, UMC Utrecht, the Netherlands

**39** Department of Cardiology, Division Heart and Lungs, University Medical Center Utrecht, Utrecht, The Netherlands

**40** Julius Center for Health Sciences and Primary Care, University Medical Center Utrecht, Utrecht, The Netherlands

**41** Department of Neurology & Alzheimer Center, VU University Medical Center, Amsterdam, The Netherlands

**42** Neurochemistry Laboratory, Clinical Chemistry Department, Amsterdam University Medical Center, Amsterdam Neuroscience, The Netherlands

**43** Department of Epidemiology and Biostatistics, VU University Medical Center, Amsterdam, The Netherlands

**44** SURFsara, Amsterdam, the Netherlands

**45** Sequence Analysis Support Core, Leiden University Medical Center, Leiden, the Netherlands

**46** University of Groningen, University Medical Center Groningen, Genomics Coordination Center, Groningen, the Netherlands

**47** Leiden Computational Biology Center, Leiden University Medical Center, Leiden, the Netherlands

**48** The Delft Bioinformatics Lab, Delft University of Technology, Delft, the Netherlands

## Notes

### Competing Interest Statement

The authors have declared no competing interest.

